# Community composition exceeds area as a predictor of long-term conservation value

**DOI:** 10.1101/2021.07.09.451808

**Authors:** Jacob D. O’Sullivan, J. Christopher D. Terry, Ramesh Wilson, Axel G. Rossberg

## Abstract

Conserving biodiversity often requires deciding which sites to prioritise for protection. Predicting the impact of habitat loss is a major challenge, however, since impacts can be distant from the perturbation in both space and time. Here we study the long-term impacts of habitat loss in a mechanistic metacommunity model in terms of both immediate extinctions and secondary species losses. We find that biomass-at-site, closely related to site area, is a poor predictor of long-term regional species losses following site removal. Knowledge of the compositional distinctness (average between-site Bray-Curtis dissimilarity) of the removed community can markedly improve the prediction of impacts at the regional scale, even when biotic responses play out at substantial spatial or temporal distance from the removed site. Fitting our model to empirical species-by-site tables describing Andean diatoms and Brazilian lichen-fungi, we show that compositional distinctness surpasses area as a predictor of long-term species losses in the empirically relevant parameter range. Our results robustly demonstrate that site area alone is not sufficient to gauge conservation priorities; analysis of compositional distinctness permits improved prioritisation at low cost.

## 1 Introduction

Habitat loss due to conversion of natural landscapes is the leading cause of biodiversity loss today (Millenium Ecosystem assessment, Reid et al. 2005; IPBES, Díaz et al. 2019). Immediate species losses that result when the habitats of endemic species are destroyed only represent part of the impact of land conversion. Additional losses may occur as ‘extinction debts’ are paid (Tilman et al., 1994). These additional extinctions can arise due to a suite of complex processes that ripple through the wider landscape, often involving multiple species (Swift and Hannon, 2010; Mouquet et al., 2011; Nee and May, 1992; Livingston et al., 2012; Terborgh et al., 2001; Taheri et al., 2021). The complexity of these ecological responses to habitat loss makes predicting long-term impacts a major challenge. Essential to meeting this challenge is an understanding of how changes in the abiotic and biotic structure of the landscape are likely to interact (Chase et al., 2020).

Decisions in conservation ecology often require identification of the least worst outcomes of landscape conversion (Wilson et al., 2009). For such assessments, sufficient mechanistic understanding of the biodiversity impacts of habitat loss is required (Figueiredo et al., 2019). Predictions of such impacts often rely on phenomenological models of species-area relationships (SAR) (Kuussaari et al., 2009; Figueiredo et al., 2019; Thompson et al., 2019), which assume that impacts follow simple scaling relationships (e.g. Thomas et al. 2004; Foster et al. 2013). However, the scaling of diversity with area arises due the fact that ecological assemblages are internally heterogeneous: it is usually not area *per se* that determines species richness but the diversity of ecological associations that a landscape can support. As such, it is plausible that metrics that directly quantify the internal diversity structure of a assemblages may outperform area alone as predictors of a site’s contribution to regional diversity.

Here, we explore the long-term outcomes of habitat destruction using a spatially explicit simulation model called the Lotka-Volterra Metacommunity Model (LVMCM), which has been shown to reproduce a large variety of well-documented patterns of spatio-temporal macroecology (O’Sullivan et al., 2019, 2021a). We model habitat conversion as the complete removal of sites from a metacommunity, and find that indeed the biotic impacts following site removal can be complex. Secondary extinctions including extinction cascades are common. These cascades can cause extinctions of populations distant from the removed site. The area of the removed site only weakly correlates with *conservation value*, which we defined as the long-term, landscape-level species loss following site removal. A stronger predictor of regional extinctions is often the compositional distinctness of the removed community, despite the cascading, far-reaching impacts removal can have. To test whether empirical systems fall into the parameter space where compositional distinctness is a stronger predictor than area, we developed a method to fit the LVMCM to biodiversity patterns derived from empirical species-by-site tables. Using this method, we demonstrate not only the higher predictive power of compositional distinctness for empirical systems, but also the wider potential of mechanistic metacommunity models as decision support tools in applied conservation (Chase et al., 2020).

### 1.1 Model description - Overview

The LVMCM extends the conventional Lotka-Volterra competition model to a spatially explicit system of connected sites (Fig. 1). It models the dynamics of a metacommunity formed by a guild of competing species. Sites differ in their suitability for each species and their total area, and nearby communities are connected by dispersal. The underlying environment is modelled as a spatially autocorrelated random field with mean zero and unit standard deviation and species responses to the environment are modelled by quadratic environmental response functions, with each species allocated a random environmental optimum. Species disperse between sites according to an exponential dispersal function. We kept the width of the environmental response function, *w*, and the characteristic dispersal length *ℓ* fixed for all species in a given simulation, but varied them systematically between simulations. For this study we kept the number of sites fixed at 20 while, under variation of these key parameters, the regional *γ*-diversity ranged from 250 to 2500 species.

**Figure 1:**
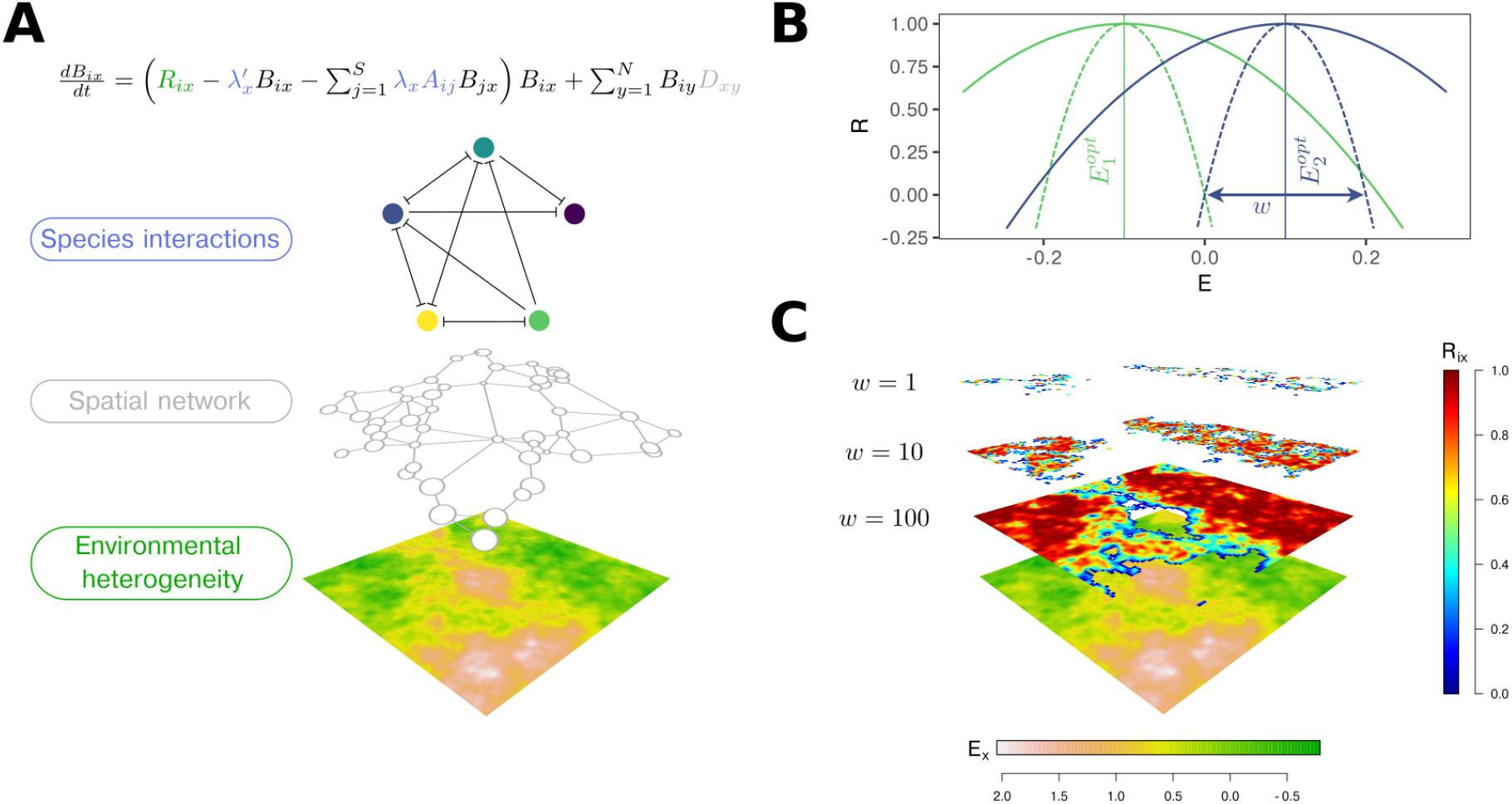
Elements of the metacommunity model. **A**: Model metacommunities include a spatially autocorrelated environment of at least one abiotic variable and a spatial network, in which local sites of unequal area are modelled by scaling the local interaction coefficients by *λ*_*x*_ and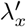. **B**: Intrinsic growth rates *R* = *R*_*ix*_, representing species’ Hutchinsonian niches, are modelled as quadratic functions of the environmental variables. Niche width is controlled by a parameter *w* and each species is assigned a randomly sampled environmental optimum 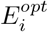 (solid lines exemplify large *w*, dashed lines small *w*, colours represent different species). **C**: Three examples illustrating of how niche width *w* affects the distribution of areas of positive *R* over the landscape, highlighting the relationship between niche width and effective heterogeneity.

**Figure 2:**
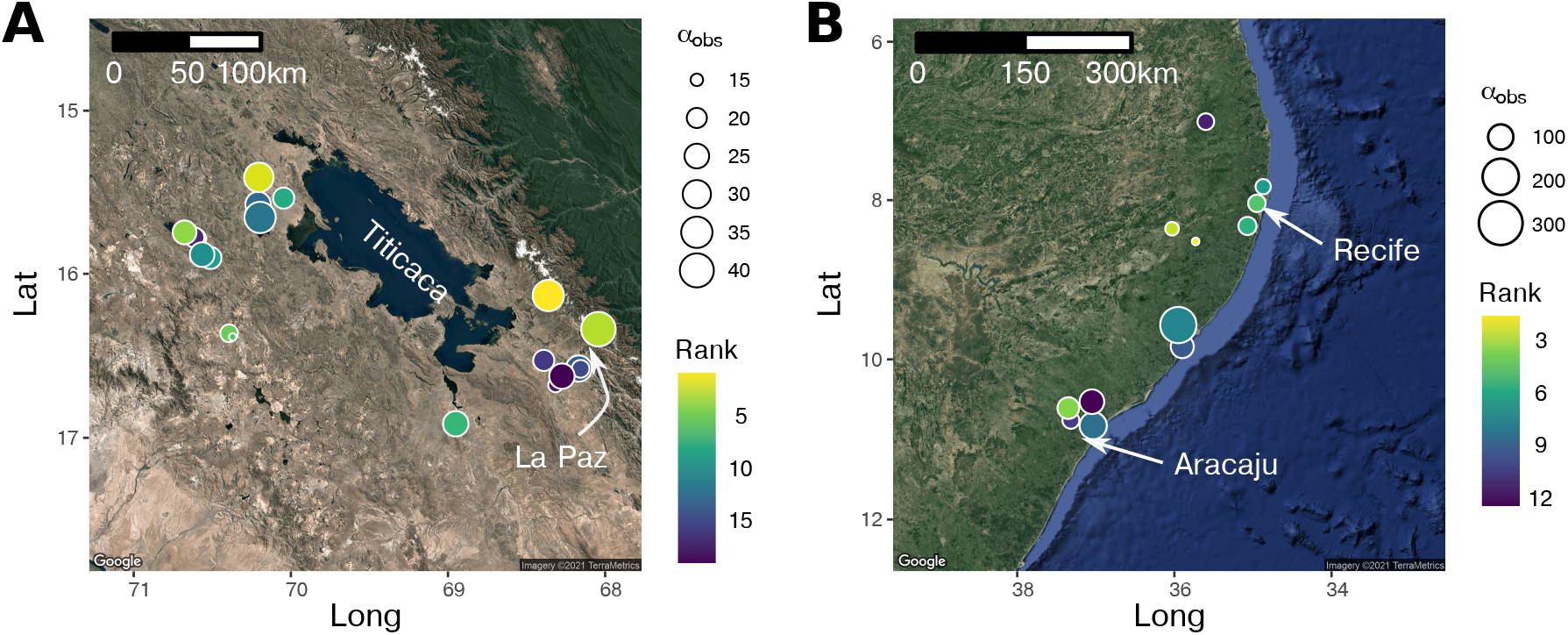
Datasets used to fit the LVMCM. **A**: The subset of the Andean diatom database (Benito et al., 2018) straddling Lake Titicaca and the Peruvian-Bolivian border. **B**: The Brazilian lichen-fungi database (da Silva Cáceres et al., 2017) with nearby samples pooled together. The colour of the points represents the ranked impact of site removal on regional diversity in *in silico* removal experiments, with the lightest colour corresponding to the site with the greatest impact. The size of the points represents the observed species richness.

We allowed metacommunity models to self-organise via a regional assembly process (Post and Pimm, 1983) through which species invade the metacommunity and distribute across the landscape, responding to the abiotic and biotic conditions in the sites. We then systematically perturbed assembled metacommunities by removing each site in turn, simulating the biotic response and measuring long-term impacts of single site removal at the regional scale (for technical details see methods).

### 1.2 Predicting long-term species losses

Process based metacommunity models like the LVMCM permit direct comparison between the immediate effects and long-term outcomes of perturbations. Immediate species losses can be predicted by asking which species have a global range limited to the removed site. We denote the predicted immediate species loss Δ*γ*_0_. In contrast, we denote by Δ*γ* the actual long-term species loss determined by simulating metacommunity dynamics. We define the conservation value of the removed site as the proportional long-term species loss, relative to the pre-disturbance regional species richness *γ*^*^.

We first we asked how Δ*γ/γ*^*^ depends on model parameters. Determining these parameters empirically, however, can be costly. Therefore we also tested how well Δ*γ/γ*^*^ can be predicted by the proportion of biomass immediately removed, 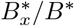, where 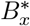 and *B*^*^ represent the pre-disturbance biomass of the removed site *x* and of the entire metacommunity, respectively. We use 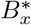 as a proxy for the area of site *x*, with the advantage that 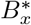 is directly represented in LVMCM model communities. As a second predictor we use a measure of compositional distinctness, 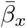, defined as the mean Bray-Curtis Bray and Curtis (1957) dissimilarity of the focal site *x* to all other sites.

We assessed the effects of 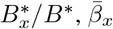and various spatial network properties (centrality measures) on Δ*γ/γ*^*^ using simple multivariate linear regression, with the best model being selected by comparison of AIC (**?**). Goodness-of-fit of predictive models was assessed using the adjusted *R*^2^. The proportions of variance explained by area and compositional distinctness were then partitioned using partial regression redundancy analysis.

### 1.3 Fitting the LVMCM to empirical data

We demonstrate how the LVMCM can be fit to data using two datasets, one describing the distributions of diatoms (D) in lakes straddling the Peruvian-Bolivian border (Benito et al., 2018), the other the distribution of Lichen-Fungi (L) in Eastern Brazil (da Silva Cáceres et al., 2017). These datasets were chosen for their high species richness and because we can assume that the ecological interaction networks of these guilds are reasonably represented by competition within a single trophic level. For dataset D, which covers a large region, we selected a subset of the full database for which the first two principal components of the key environmental variables formed a distinct cluster, including 19 sites and 221 taxa. For dataset L we reduced the total number of sites (and the computational effort) by pooling observations separated by less than 20km. This left 12 sites and 784 taxa.

In the Appendix we give a detailed summary of the model fitting procedure that we outline here. The spatial structure (spatial distance matrix) and key environmental variables from the empirical dataset were input directly into the model, defining the distance between sites and the underlying abiotic environment. We then used a battery of simulations to estimate the abiotic niche width parameter *w* that best fit the empirical species by site table. This was done by quantitatively comparing the occupancy frequency distribution (OFD) (McGeoch and Gaston, 2002) for the empirical observation to that of the simulation under systematic variation of *w*.

The LVMCM is a parameter rich model, making direct fitting to data impossible. Therefore we developed a simplified patch occupancy model that predicts high-level metacommunity properties with a single fitting parameter, which we termed the ‘mixing rate’ *m*. Elsewhere we have shown that this simple framework can explain the spatial and temporal properties of riverine communities (O’Sullivan et al., 2021b). While unsuitable for in-depth analyses of biotic responses to perturbation, the simplicity of the patch occupancy model means it can be directly fitted to both empirical observations and LVMCM model communities, thus serving as a bridge between the complex LVMCM simulations and empirical ecosystems. The theory behind this simple model and the procedure for model fitting are summarised in the Appendix.

The parameter *m* can be estimated from the OFD for a given dataset. For LVMCM models with spatial and environmental distributions matching the observation, the abiotic niche width parameter space of the model was scanned. The value of *w* that best reproduces the mixing rate of the empirical observation was then located. This value gives a meaningful estimate of the typical niche widths relative to the total environmental variation in the assemblage which includes explicit consideration of the role of biotic interactions in determining species ranges.

## 2 Results and discussion

Even though none of the sites removed in simulation experiments comprised more than 12% of the total regional biomass/area, we detected regional extinction of at least one species in over 75% of cases. The highest impact of a single site removal was a proportional loss of Δ*γ*/*γ*^*^ = 0.23. In a supplementary movie we show examples of extinction cascades following site removal, demonstrating species loss from sites both near and distant from the removed site.

### 2.1 The dynamics of extinction following habitat removal

Proportional long-term species loss Δ*γ*/*γ*^*^ decreased with increasing abiotic niche width (Fig. 3A). This is plausible, since with wider niches single-site removal tends to remove a smaller proportion of the area representing a species’ Hutchinsonian niche. By contrast we found that dispersal length had surprisingly little effect on Δ*γ*/*γ*^*^ (Fig. 3A).

**Figure 3:**
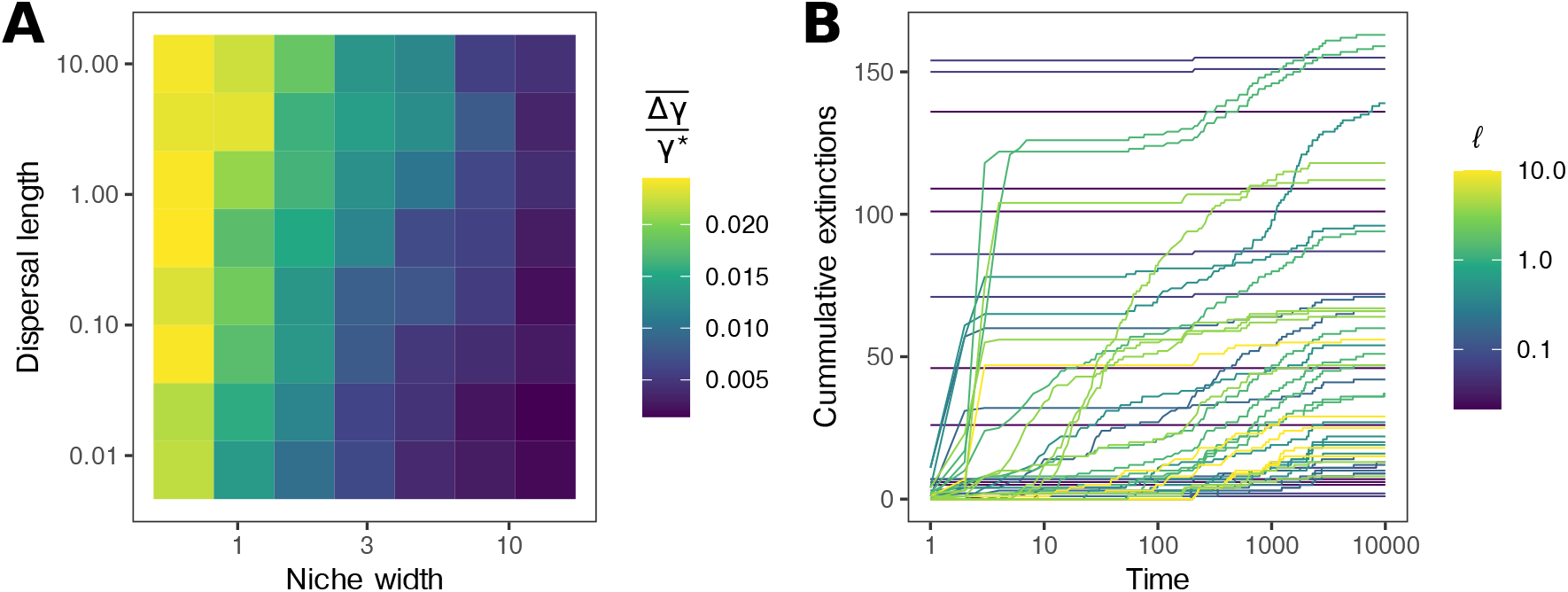
Species extinctions following site removal. **A**: The average proportional species losses for all combinations of the abiotic niche width and dispersal length. **B**: Secondary extinctions occur due to the interuption of source-sink dynamics and extinction cascades. Here we show the outcome of removing sites for the smallest niche width *w* = 0.63.

The process by which species are lost following site removals was complex. In Fig. 3B we show a random sample of these complex extinction dynamics for various dispersal lengths with abiotic niche widths fixed at *w* = 0.63, the value for which impacts are largest. We find that site removals can either lead to the loss of endemics only (cummulative extinctions effectively independent of time lag), or they can lead to extinction cascades which can play out over long times. These extinction cascades, particularly prevalent in higher dispersing metacommunities, demonstrate the complexity of potential metacommunity responses to site removal.

The secondary extinctions of non-endemics can be categorised into two distinct types. Those occurring due the disruption of mass effects (sink populations which are lost following the removal of source sites), and those occurring due to a complex restructuring of the metacommunity as species ranges shift in space. Extinctions of the first type typically occur within around 50 unit times and in sites adjacent to the removed site. The average distance between the site in which a species was last detected and the original perturbation was 0.05*L*, where *L* is the side length of the model landscape, in the first 50 unit times (smallest niche width, across dispersal lengths). The second type can occur much later and at almost any site in the metacommunity. For extinctions occurring after more than 50 unit times, the average distance of the final local extinction in a given species’ decline to extinction was 0.45*L*. For species lost after 120 unit times, the distance from the initial perturbation had a mean (0.5004*L*) that was statistically indistinguishable from the average inter-site distance (0.5002*L*), suggesting the location of subsequent species losses was essentially random with respect to the initial impact.

### 2.2 Predicting long-term impacts based on empirically measurable quantities

Predicting the outcome of single site removals, given the complexity of the biotic response (Fig.3C) is non-trivial. It is not the case that species losses in the model can be generically explained by reference to immediate losses of endemics. Instead, we test whether area and compositional distinctness, both empirically accessible properties of communities, correlate with the long-term outcomes of the often complex structural redistribution precipitated by single site removal.

We find a clear positive association between the conservation value of sites and 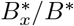, but with substantial spread (Fig. 4, Spearman’s *ρ* = 0.33, all parameter combinations pooled). In contrast, long-term species losses were more strongly associated with compositional distinctness of the removed site 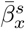(Fig. 4, *ρ* = 0.52, all parameter combinations pooled).

**Figure 4:**
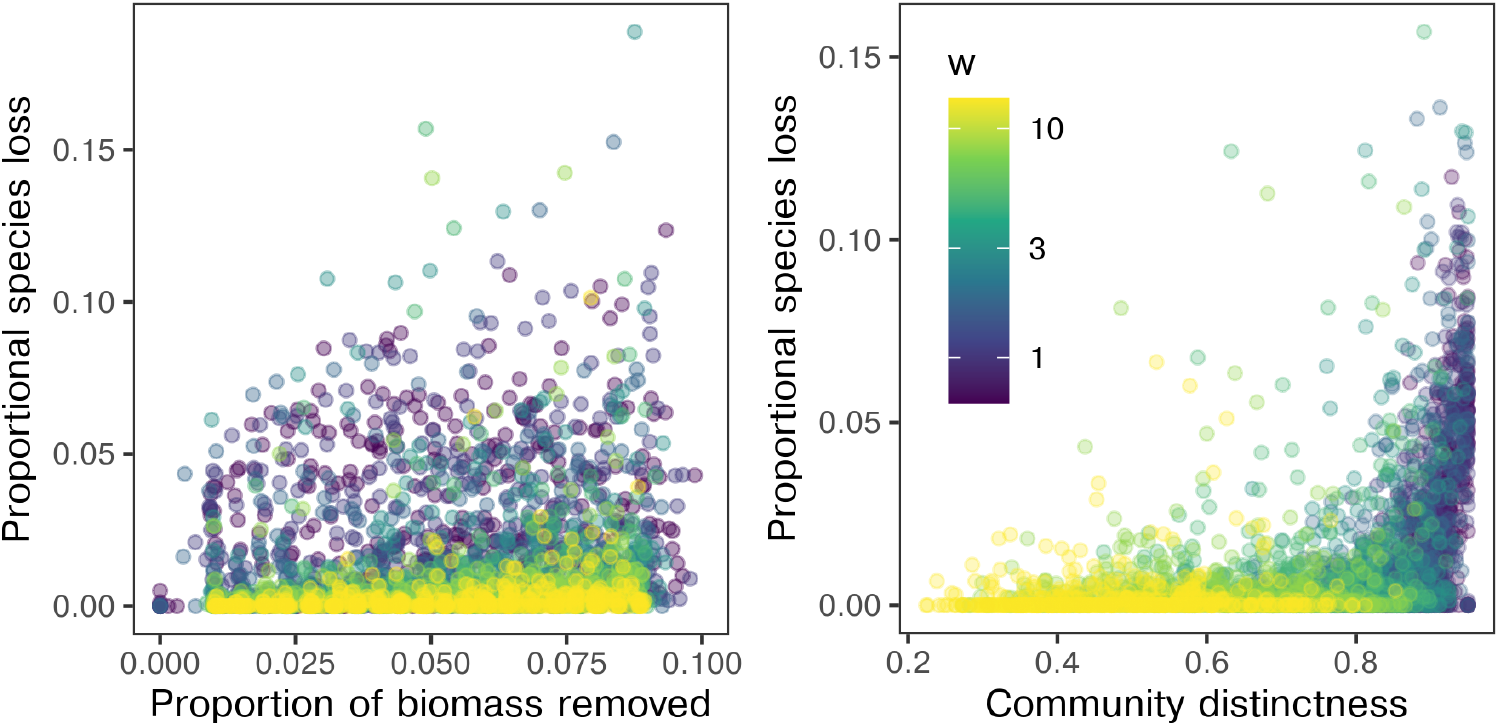
Predicting long-term species losses. The proportional long-term species losses following site removal plotted against the proportion of biomass removed initially, a proxy for site area, and the compositional distinctness of the removed site. Here we show a random subset of the simulated removal experiments for visual clarity.

The best model selected by AIC included both proportional area removed and compositional distinctness of removed site as predictors of long-term species losses. Remarkably, none of the standard centrality measures available (degree-, closeness-, betweenness-, eigenvector centrality) were selected as predictors.

Decomposing the simulation results by *w*, we find that the strength of the association between long-term impacts and the properties of the removed site varies considerably with nice width. In Fig. 5 we show how the *R*^2^ and the regression coefficients of the multivariate linear models vary over the parameter space.

**Figure 5:**
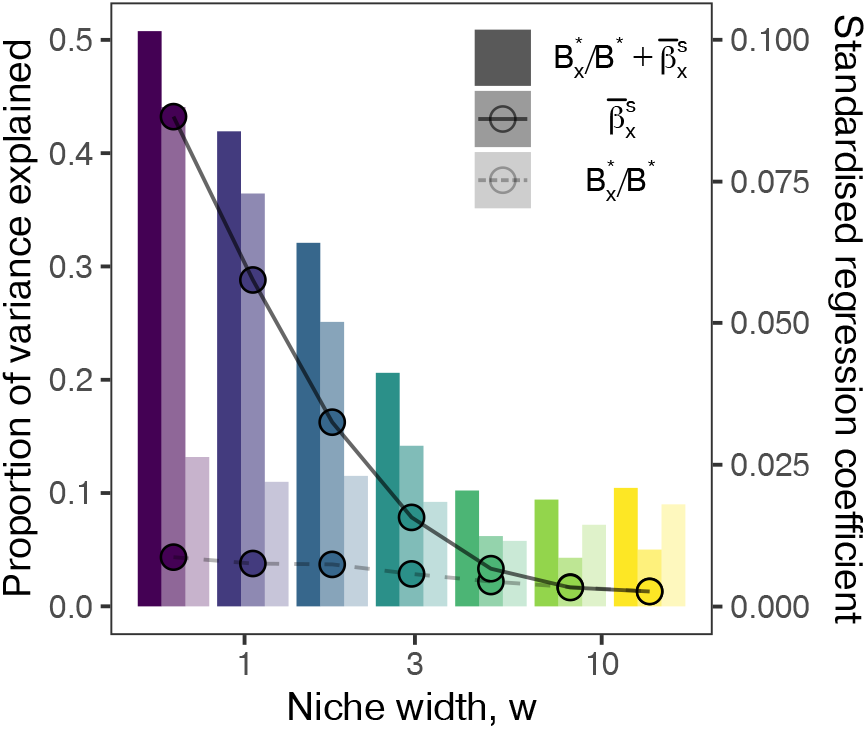
Dependence of predictive power on niche width. We find a clear decay with increasing niche width *w* in the variance explained (bars) by both the full model (combining both predictors) and model based on distinctness. In contrast, the proportion of variance explained by area was largely independent of *w*. Coincident with the decay in variance explained, the standardised regression coefficients (points) also decayed with niche width. The standardised effects of area (dashed line) was consistently lower than that of compositional distinctness (solid line).

For the smallest niche widths studied here, the *R*^2^ (bars in Fig. 5) of the full model (both predictors) increased up to 0.55 for the smallest niche width. For large niche widths *R*^2^ stabilised at 0.13. Thus, predicting Δ*γ*/*γ*^*^ when niche widths are large relative to environmental variation is particularly challenging. This is partly due to the fact that long-term species losses are rare when environments are benign, but may also reflect the fact that the biotic response to perturbation depends much more on dispersal and local competitive effects–absent from the multivariate regression–where abiotic filtering is largely neutral.

The proportion of variance explained by area following partial regression redundancy analysis was largely consistent across niche width parameterisations. In contrast, the proportion of variance explained by compositional distinctness decayed with increasing niche width. As a result, for the largest niche widths studied, the variance explained by area actually exceeds that by composition, though only once total *R*^2^ had dropped to its minimum. The standardised regression coefficients (points in Fig. 5) show that for all but the widest abiotic niches, the effect of compositional distinctness exceeded that of area on long-term losses (solid and dashed lines respectively).

By repeating this analysis for random sub-sets of the simulated metacommunities containing a fixed number of species, we verified that these differences are not due to differences in overall species richness of low- and high-*w* metacommunities.

Thus, we conclude that compositional distinctness typically outperforms biomass as a predictor of long-term losses, despite the fact that the population scale impacts are not uniquely felt in or near the impacted site. In order to estimate the long-term effects of habitat destruction on biodiversity, it is critically important to take the community composition of the affected areas into account.

### 2.3 Fitting the LVMCM to empirical observation

In view of the strong parameter dependence of the predictive power of compositional distinctness, it is critical to ascertain which region of the parameter space is most representative of natural ecosystems at large scales. We constructed a set of LVMCM metacommunities with spatial and environmental distributions taken from the two empirical datasets. By comparing the mixing rate *m* estimated from the empirical species by site table to that for LVMCM models we found that dataset D was best fit when *w* = 3.46. In the case of dataset L *w* = 1.08 gave the closest match to observations.

The simulations deviate from the empirical data in two of important respects (Fig. 6A). The empirical OFDs tend to have a sharper peaks at single site occupancy than the simulations and slightly fatter right tails. This is most likely due to the model’s simplifying assumption that all species have the same abiotic niche width. Inspection of the OFD for various niche widths (Fig. S4) suggests that a better fit to the empirical distribution could be achieved by relaxing this assumption. We did not attempt to incorporate a distribution in environmental specialism in our model to avoid over-parameterization and because the shape of such distributions is not well understood empirically (O’Sullivan et al., 2021b).

**Figure 6:**
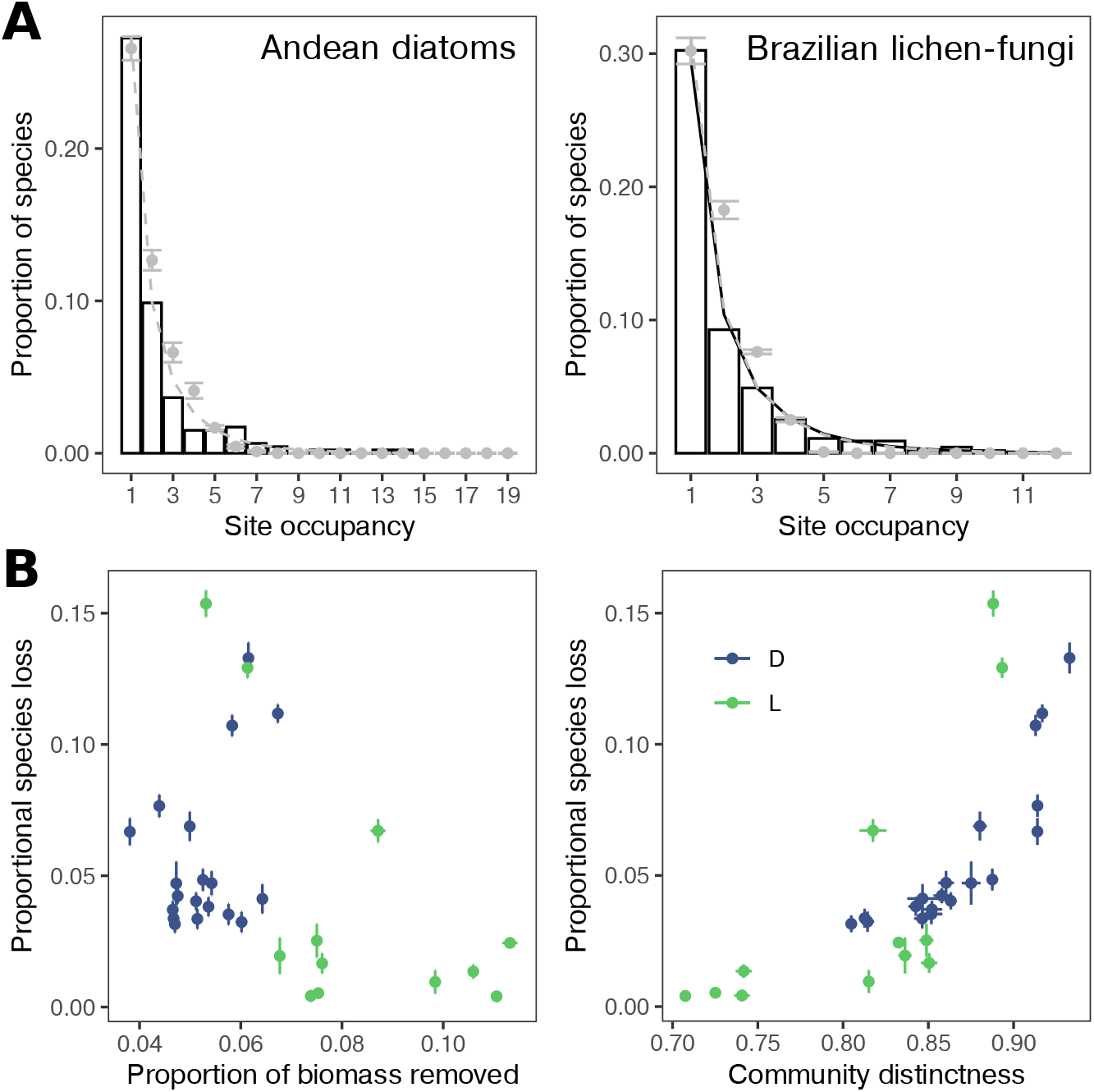
Fitting the LVMCM to empirical OFD and simulated habitat loss. **A**: Bars represent the empirical OFDs The black curves are the OFD of the patch dynamics model with the mixing rate fitted to the observation. Grey points are the mean occupancies of 10 replicate LVMCM simulations with parameters that best fit the empirical observations. The grey dashed lines are the OFDs of the patch dynamics model fitted to the simulation. **B**: The outcomes of the simulated removal experiments for the best fitting parameter combination. Each point represents a site, error bars are the standard deviation in the outcome over 10 replicate assemblies.

Despite these differences, our simple fitting procedure generated model metacommunities with macroecological spatial structure (OFD) approximately matching that of the empirical observation. The key finding is that both datasets are fit by abiotic niche widths of order *w ≈* 1—representing around one standard deviation of the landscape scale heterogeneity—suggesting the parameter space in which compositional dissimilarity substantially exceeds biomass as a predictor of conservation value is the most biologically relevant.

Using these fitted models, we also performed a direct mechanistic assessment of the rank conservation priority of each site (Fig. 2), performing systematic removal experiments as described above. In these experiments the proportional drop in diversity Δ*γ*/*γ*^*^ after removal of a single site ranged from 0.03 to 0.13 for dataset D and 0.004 to 0.15% for dataset L (Fig. 6B). In both cases, 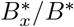was a rather poor predictor of simulated species losses. On the other hand, a strong non-linear relationship between long-term impact and 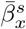was found, consistent with the results shown in Fig. 5. For dataset D, conservation priority typically increased with site altitude as spatial isolation and deviation from environmental averages increased. Surprisingly, for dataset L the greatest impacts occurred when either of the two smallest sites were removed. This apparently incongruous result is explained by the fact that for this dataset observed species richness was negatively associated with environmental rarity, measured as the mean Euclidean distance of focal site from all other sites in environment space (Spearman’s *ρ* = *−*0.56, *p* = 0.057). In dataset L, the fact that small sites occupy the most distinct environments, and therefore host the most distinct communities, means the relationship between long-term impact and site area was reversed compared to SAR-based expectation.

We acknowledge that testing the accuracy of these prediction is a challenge. The present study demonstrates the possibility of a bespoke, simulation-based assessment of conservation value, but leaves further refinements and validation of the procedure to future work.

## 3 Conclusions

The biotic response of metacommunities to localised perturbation can be complex and far reaching. While endemic species are by definition lost immediately, secondary extinctions can be substantial as metacommunities restructure. Predicting the long-term impacts of this restructuring is a major challenge, and one that currently can only be studied using process-based metacommunity models like the LVMCM. With this study we have shown that compositional distinctness can substantially exceed area alone as a predictor of long-term impacts, including secondary extinctions, on biodiversity following habitat destruction.

Compositional distinctness, measured in terms of average dissimilarity, is likely to be empirically accessible because Bray-Curtis dissimilarity between sites is numerically dominated by the locally dominatant populations, the sizes of which are readily quantified. However, the usefulness of the predictor is based not only to what it tells us about the dominating species, but also what the dominating set of species tells us about the abiotic characteristics of a site. The compositional distinctness of the dominant taxa is likely to be informative as to the distinctness of the non-dominant taxa, which is harder to measure.

While at first sight it might appear that application of this predictor requires a well-defined metacommunity to average over (which in practice may be ambiguous), this is not actually the case. Addition or omission of sites from a dataset affects ordering of two sites in terms of mean Bray-Curtis dissimilarity only for those other sites in the dataset that have compositions similar to either of the two sites.

To illustrate the relevance of our results, we note that the recently published first draft of the Post-2020 Global Biodiversity Framework^1^ sets as a its primary goal the enhancement of ecosystem integrity, including increasing the area and connectivity of natural ecosystems, ensuring the robustness of populations and the reduction in extinction rates. Accompanying this document is a set of indicators proposed to help monitor progress toward these goals^2^. We can roughly categorise the 56 indicators applicable to the primary goal of the Framework into those relating to (i) area, extent or coverage, (ii) species-level assessments (of extinction risk, habitat integretity etc.) (iii) intactness (describing the proportion of historical assemblages that persists today) and (iv) spatial beta diversity intrinsic to an ecological unit (region, nation). The number of indicators in each category in the current draft are (i) 19, (ii) 12, (iii) 3 and (iv) 2. Thus, while our demonstration that compositional rarity is a key biotic quantity that must be conserved in order to protect regional biotas may be intuitive in hindsight, the weighting of area and composition in key management tools currently remains strongly skewed toward area-based measures. We therefore advocate for increased prioritisation of descriptors of the internal structure of biodiversity when predicting impacts and setting conservation targets.

## 4 Materials and methods

### 4.1 Model description - Environmental heterogeneity and spatial structure

Environmental heterogeneity (Fig. 1A) was modelled by assigning to each each site *x* a set of *K* independent random variables *E*_*kx*_ (1 *≤ k ≤ K*), representing, e.g. the principal component(s) of abiotic environmental variation. The *E*_*kx*_ were sampled from spatially correlated Gaussian random fields (Adler, 2010) with mean 0, standard deviation 1, and correlation length 1.

Each species *i* in the model was allocated environmental optima 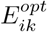 for each environmental variable *k*, sampled from uniform distributions in the range [1.25 · min_*x*_ *E*_*xk*_; 1.25 · max_*x*_ *E*_*xk*_]. The effective heterogeneity of the environment was controlled by varying the niche width parameter *w*. The intrinsic growth rate of the *i*th species in site *x* was computed as

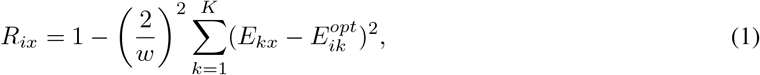

that is, to simplify parameterisation, we assumed identical niche widths for all species and all *n* independent components of environmental variation. For the random metacommunities we set *n* to either 1 or 2 and observed no major change in outcomes. For the fitted metacommunites, the first two principal components of the observed environmental variation were used.

The spatial network of sites (Fig. 1A) was modelled by randomly sampling Cartesian coordinates from a uniform distribution in the range 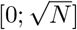 where *N* is the number of sites. As in previous studies (O’Sullivan et al., 2019, 2021a), we linked nearby sites using a Gabriel graph (Gabriel and Sokal, 1969).

### 4.2 Model description - Scaling local site area

An essential technical innovation that made this study possible is a technique we developed to model local differences in site area by scaling the intensity of local ecological interactions. The precise functional form of the area dependence of intra- and inter-specific ecological interactions is not currently known. In order to overcome this knowledge gap we assembled metacommunities of different total area *a* and measured the decay in effective interaction strengths. Previous work has shown that regional scale interaction coefficients *C*_*ij*_ describing the interaction between species *i* and *j* can be estimated for models by perturbing the biomass of each species in turn and summing the impacts on all other species over the metacommunity (O’Sullivan et al. 2019 and S1 of the appendix for details). The resulting interaction matrix **C** captures the combined effects of differences in environmental preference, limited dispersal, indirect and direct interactions.

The decay in average interaction strength with area, which we refer to as the *competition area relation* (CAR), can be modelled by two power laws, one for inter-specific interactions, 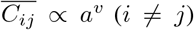, and one for intraspecific interactions, 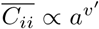 (Fig.S1A). The exponents *v* and *v*′ depend on model parameters, in particular the abiotic niche width *w*, which strongly affects species’ range sizes (we found the effect of dispersal length to be weak by comparison). The CAR can be incorporated into the model dynamics in order to model variation in the local biomass (area) between sites by scaling the site-specific interaction matrix **A**_*x*_. Thus we model each site as an implicit sub-network of various nodes and scale the interactions to those expected for the corresponding (sub-)metacommunity.

Formally, if **A**_0_ is a hollow matrix with zeros on the diagonal, the scaled matrix is given by

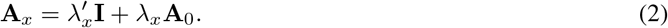

where **I** is the identity matrix, and

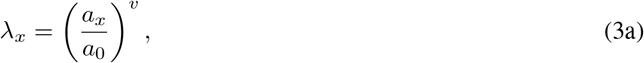

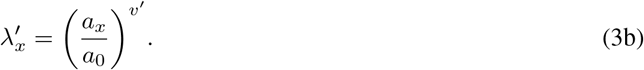

Here *a*_*x*_ is the area of the *x*th site and *a*_0_ the area of a reference site with (unscaled) interaction matrix **A** = **I** +**A**_0_. We measure site area in units such that *a*_0_ = 1. For random metacommunities, the *a*_*x*_ values were randomly assigned to sites *x* such that they covered the range from 1 to 30 biomass units in equal intervals. For fitted metacommunities, the *a*_*x*_ were extracted from the empirical species by site tables (see below). The exponents *v* and *v*′ were set based on the relationships between the CAR and SAR found in simulations, assuming that within-site SARs are typically linear (see S1 of the appendix).

### 4.3 Model description - Metacommunity dynamics

Metacommunity dynamics were modelled using a spatial extension to the classic Lotka-Volterra community model (O’Sullivan et al., 2019, 2021a)

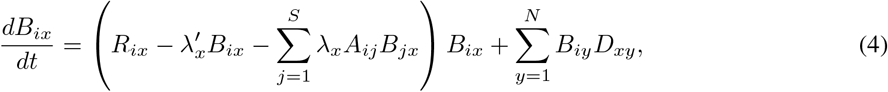

where *B*_*ix*_ represents the biomass of the *i*th species in the *x*th site. *R*_*ix*_ are the intrinsic growth rates, which vary across the landscape.

For simplicity, the unscaled local interspecific interaction coefficients *A*_*ij*_ were sampled from a discrete distribution with *P* (*A*_*ij*_ = 0.3) = 0.3, and *A*_*ij*_ = 0 otherwise. *D*_*xy*_ are the elements of the spatial connectivity matrix describing the inter-site dispersal. Emigration and immigration rates are given by *D*_*xx*_ = −*e, D*_*xy*_ = (*e/k*_*y*_) exp(*−d*_*xy*_*/l*) for sites *x, y* connected by the spatial network, and *D*_*xy*_ = 0 otherwise; *e* is an emigration rate, *d*_*xy*_ the distance between sites *x, y*, and *ℓ* the characteristic dispersal length, which was systematically varied. The normalisation *k*_*y*_ represents the unweighted degree of the *y*th site.

### 4.4 Model description - Metacommunity assembly and removal experiments

Model metacommunities were assembled by iteratively generating species *i* with randomly sampled 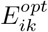 and *A*_*ij*_, and introducing them as invaders to the model at low biomass. Dynamics were then simulated over 500 unit times using Eq. 4. Before each invasion, the metacommunity was scanned for any species whose biomass had fallen below the extinction threshold of 10^−4^ biomass units in all sites. These species were considered regionally extinct and removed from the model. Metacommunity models assembled in this fashion eventually saturated with respect to both average local and regional species richness (O’Sullivan et al., 2019) due to the onset of ecological structural instability (Rossberg, 2013). After saturation is reached, each invasion generates, on average, a single extinction.

Following pilot studies metacommunities were assembled with *w* assigned 10 values logarithmically spaced in the range 0.5 *≤ w ≤* 15. The parameter *ℓ* was logarithmically spaced in the range 2 · 10^−2^ *≤ w ≤* 10, again with 10 distinct values included. In both cases, a couple extreme values were excluded from the analysis either because beyond a threshold, no further change in outcomes was observed or, in the case of *ℓ*, because very small values lead to numerical errors in the simulation.

After assembling model metacommunities of 20 sites to regional diversity limits, each site in turn, and all associated edges were systematically removed and the simulation advanced over 10^4^ unit times. For completeness, the dispersal matrix **D** was updated to reflect the new degree distributions.

## Appendix

### S1 Harvesting experiment and the CAR

Following O’Sullivan et al. (2019) we computed an *S × S* spatially unresolved competition matrix **C** that summarised biotic interactions at the regional scale. For this we constructed a spatially unresolved Lotka-Volterra system,

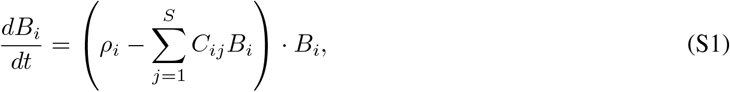

where *B*_*i*_ represents total biomass of species *i*, as an approximation of the spatially resolved model. The regional scale interaction coefficients *C*_*ij*_ were then estimated using a computational harvesting experiment, which assumes that interaction strengths can be inferred from the changes in regional abundances that result from controlled changes in the regional abundances of harvested species (Gilbert et al., 2014).

For the unresolved model, Eq. S1, the harvesting of species *i* at a rate *h*, represented by substituting *ρ*_*i*_ *− h* for *ρ*_*i*_ in Eq. S1, produces a shift in the equilibrium biomasses given by

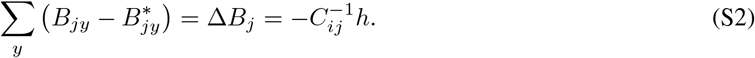

where **B**^*^ represents the unharvested equilibrium. Near an equilibrium 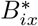of the full metacommunity model,

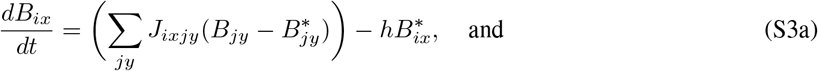

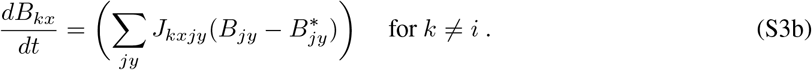

where *h* is the harvesting rate, and **J** is the numerically estimated *SN × SN* Jacobian of the full metacommunity model. We vectorize the matrix **B** (denoted 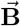) to match the dimensionality of the spatially resolved Jacobian, and the write the equilibrium condition for Eq. S3a as

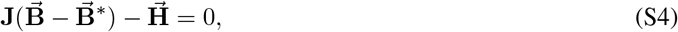

where the elements of vector 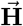 are 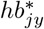 for *j* = *i* and 0 otherwise. From this, we obtain

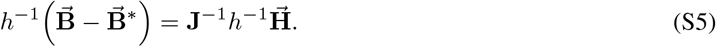

The left hand side of Eq. S5 represents the local change in biomasses due to the harvesting of the focal species *I* per unit *h*. From Eq. S5 we compute the change in total biomass 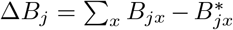 for each species *j*, which gives row *i* of **C**^*−*1^ (Eq. S2). Iterating over all species *i* = 1 … *S*, we computed **C**^*−*1^ and from this the spatially unresolved interaction matrix **C**. This method requires that the metacommunity is at equilibrium with respect to biomass. In practice, this assumption is violated by the onset of autonomous compositional turnover for large *N*. For this reason we restricted our analysis to a range of *N* for which autonomous turnover is absent or low.

The approximate functional form of the CAR and the scaling relationships between *v, v*′, *z* and *w* can be studied by assembling metacommunities of various numbers of sites (total area) and computing the mean regionalscale interaction coefficients 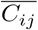 using the harvesting approach (Figs. S1A). The exponents *v* and *v*′ depend on model parameters, in particular the abiotic niche width *w*, which strongly affects species’ range sizes (we found the effect of dispersal length to be weak by comparison). Future developments in the theory of LV metacommunities are likely to reveal close relationships between the CAR, the SAR (*S ∝ a*^*z*^) and the effective heterogeneity of the landscape (here determined mainly by *w*). In lieu of analytic results, we report the statistical relationships between these key quantities in Figs. S1B-D. The exponent *z* decays with abiotic niche width *w*, suggesting greater abiotic heterogeneity (smaller niche width) results in a steeper SAR. This is consistent with previous results (O’Sullivan et al., 2019) and qualitatively matches the observation of a steep phase in the empirical SAR at the highest spatial scales (greatest effective heterogeneity) (Allen and White, 2003; Rosindell and Cornell, 2007; Storch, 2016). The ratio of the exponents of the CAR *v*′*/v ≥* 1 holds over the examined parameter space and further that *v*′*/v ≈* 1 *− z*. This suggests a pivotal role for the ratio between the area dependence of inter- and intra-specific interaction coefficients in determining the accumulation of species with area, and thereby the exponents of the SAR. Finally, the rate of decay of the intra-specific interaction coefficient, determined by *v*′, and abiotic niche width *w* are negatively correlated. When abiotic niche widths are very small, intra-specific interaction coefficients are largely insensitive to total area (*v*′ approaches 0), since species ranges are primarily limited by the environment. In contrast, when niche widths are large and species distributions can encompass much of the landscape, effective intra-specific interactions decay more rapidly with total area (*v*′ approaches *v*, Fig. S1A). The linear associations revealed in Figs. S1B-D may hold only for a subset of the parameter space of the model, however this subset includes the parameter combinations that we find to best fit empirical data.

**Figure S1:**
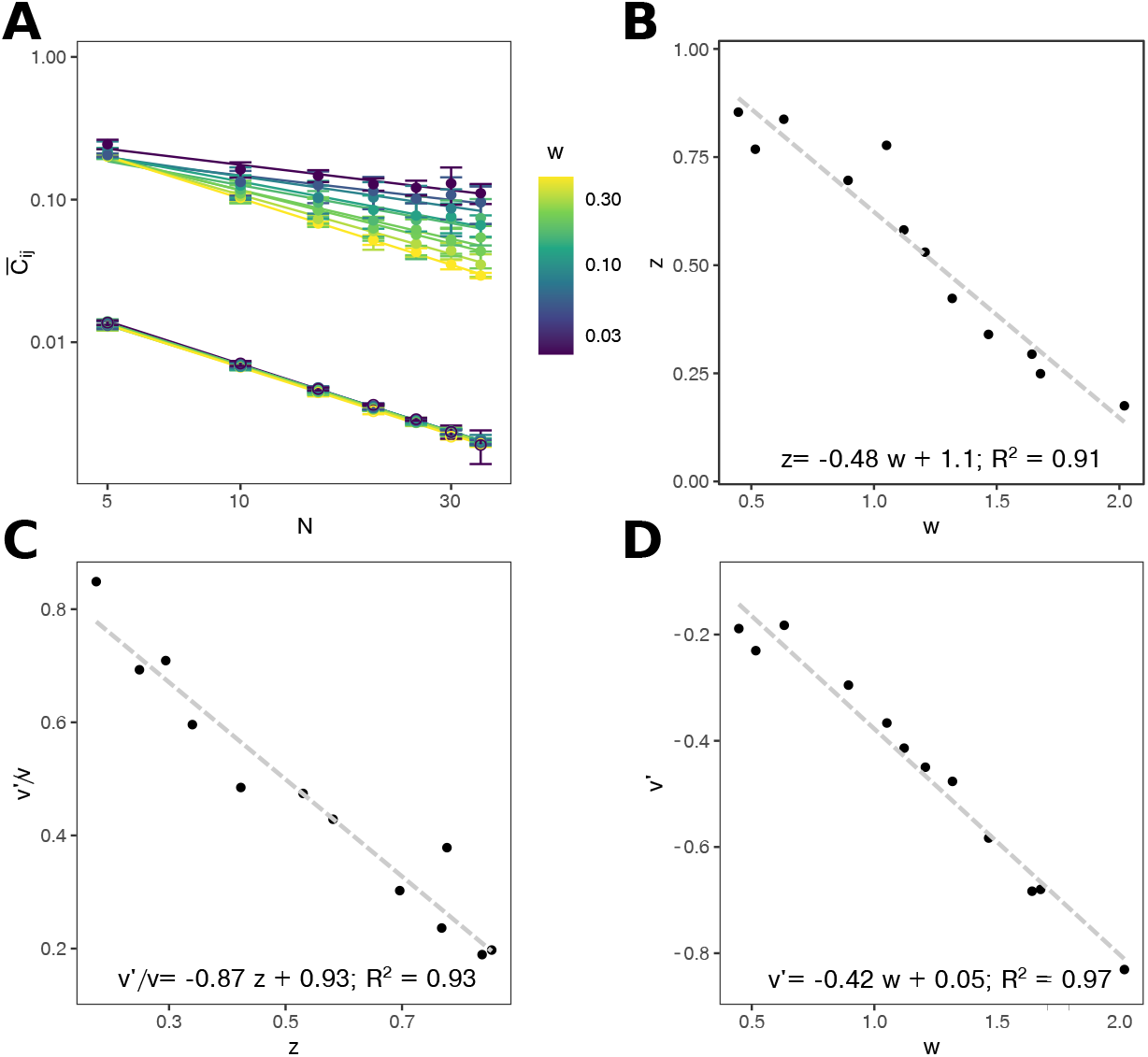
Competition area relations in the LVMCM. Assembling metacommunities with different total area reveals a power-law decay in average regional scale competitive interaction coefficients (A). We show the CAR for random metacommunities of 5 ≤ *N* ≤ 35 sites for a range of niche widths *w*. The area dependence of mean inter-specific interaction strengths are largely unimpacted by niche width. In contrast there is a substantial effect of *w* on the exponent *v*^*ℓ*^ for intra-specific interactions. Linearly regressing the exponent *z* of the species-area relation (SAR) against niche width *w* (B), *v*^*ℓ*^*/v* against *z* (C) and *v*^*ℓ*^ against *w* (D) offers some indication of the mechanisms through which effective heterogeneity impacts the SAR and inform the scaling of interactions in our model. In B-D, the exponents *z, v* and *v*^*ℓ*^ were computed by assembling metacommunities for given *w* and *N* = 5, 10, 15, 20, 25, 30, 35 sites in 10-fold replication, such that each point in B-D represents an regression across 70 independent simulations.

We assume that accumulation of diversity with area within sites is linear, that is, that *z* = 1. This simplification is biologically reasonable given the tendency for the initial phase of empirical SARs to be linear. From the relationships between the key quantities summarised in Fig. S1 we can therefore set the exponents of the CAR in this linear phase as *v* = *−*0.59, *v*^*ℓ*^ = *−*0.04.

**Figure S2:**
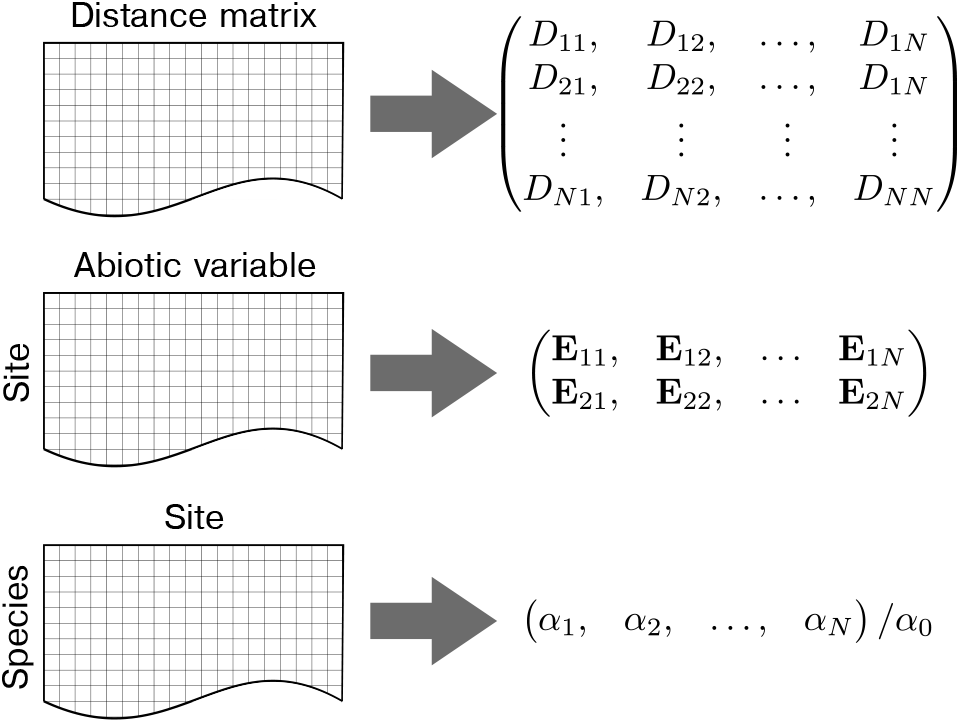
Extracting data for fitting to empirical observation. To fit the LVMCM to empirical observation we extract spatial coordinates, scaled to a square of longest side length 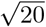 so that parameter values correspond to those used in the random metacommunity simulations. From the species by site presence-absence table, we extract the local site richness, presumed to scale with site area (quality). This is then used to generate *λ*, and *λ*^*ℓ*^ which scale the interaction coefficients **A**_*ij*_ so that the simulated metacommunity reproduces the observed distribution in *α*-diversity. From the available environmental metadata we extract the first two principle components and input these as the vectors **E**_*k*_ in the simulation.

### S2 Detailed procedure for fitting to empirical data

Our approach to fitting the LVMCM to empirical species-by-site tables is as follows: We determined for each data set the Universal Transverse Mercator coordinates of each site, the two principal components of a multivariate distribution of environmental variables scaled to mean zero and unit variance, and the species richness recorded at each site (Fig. S2). The first two are input directly into the model. For better comparison to the parameters used above, we scaled the spatial network such that it occupies an arena of which the longest extension was 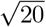. We again use a Gabriel graph to model the landscape connectivity, but note that alternative networks based on topographical data could be readily implemented.

To characterise environmental variability in dataset D, we followed the approach of the authors of the original study and used the NIPALS (nonlinear estimation by iterative partial least squares) algorithm, which allows ordination of datasets including missing values (Ibáñez et al., 2012), to extract the principal components of 14 environmental variables describing water chemistry, average climate, climate seasonality and variables describing the topography of landscape (see Benito et al. 2018 for details). Dataset L did not include quantitative environmental variables. We therefore extracted elevation, mean annual temperature, and mean annual precipitation data from publicly available remote sensing databases (further details given in the S4).

Because the ecological interactions between the observed species are insufficiently known, we randomly sampled the interaction coefficients **A**_*ij*_ from the same discrete distribution used in the random metacommunities. For a community model with no spatial extent for which *P* (*A*_*ij*_ = 0.3) = 0.3, and otherwise *A*_*ij*_ = 0, the onset of ecological structural instability occurs when *S*_0_ *≈* 40. From the SAR the effective area of a given site can be estimated using the equation

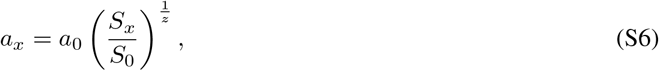

where we assume *z* = 1 as explained above. Substituting Eq. S6 into Eq. 2, the scaling of the competitive overlap matrix for the fitted metacommunity model can be written as

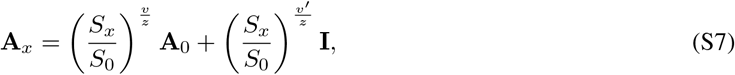

where *S*_*x*_, the observed richness at a given site, is extracted from the empirical data.

When assembling fitted metacommunity models, we again made the simplifying assumption that abiotic niche widths (*w*) are identical for both environmental variables and all species. Greater accuracy in model fitting could be achieved by estimating the typical abiotic niche widths of each variable independently, and by incorporting inter-specific variation in niche width.

For simulated LVMCM metacommunities with environmental and topographic distributions matching empirical observations we fit the parameter *w* based on the match between the observed and simulated occupancy frequency distribution (OFD) (McGeoch and Gaston, 2002), the proportion of species occupying a given number of sites (ranging from 1 for the site with the highest estimated conservation value to *N* the number of sites in the dataset). Since we find the parameter *ℓ* to have very little impact on the response to site removal, this was kept fixed at an intermediate value when fitting to observations. The OFD of the observation was quantitatively compared to the simulation by fitting to a pseudo-mechanistic patch occupancy approximation of the LVMCM, which is outlined in S3 of the appendix and will be explored in more detail in future work.

### S3 Patch dynamics fitting model

We use a simple locally saturated patch occupancy model (LSPOM) as an intermediary for comparison of the occupancy frequency distributions (OFD) of LVMCM and empirical datasets. This is advantageous because the LSPOM has only a single fitting parameter, called the ‘mixing rate’, and can be solved analytically. Thus, comparison between OFDs reduced to comparison of this single fitting parameter. We discuss motivation and validity of the LSPOM in detail elsewhere (O’Sullivan et al., 2021b). Here, we only provide a bare-bones statement of the model and its analytic solution.

The LSPOM conceptualizes a metacommunity as a spatially implicit species-by-site occupancy matrix describing an assemblage of *S* species occupying *N* patches, each capable of hosting *n* populations. This constraint on local richness reflects the fact that in each patch in the full LVMCM the number of co-existing species is limited by ecological structural stability constraints. Thus, each site can be thought to contain *n* ‘slots’, each of which can host a single local population. Each species present in the metacommunity occupies one or more slots, and the distribution of the numbers of slots occupied by species is the OFD. If most species occupy much fewer slots than there are patches (as we find here), then one can disregard, in a first instance, the constraint that a species cannot occupy more than one slot in the same patch.

At each time step of the model a new species invades the metacommunity from outside. Its initial site occupancy is set to 1, i.e. each new species colonises a single site in the metacommunity. Simultaneously, each extant local population colonises an additional site with a probability *m/*(*nN*), where *m* is the mixing rate. Each colonisation event, either from within or from outside the metacommunity, produces a corresponding extirpation event of a randomly selected occupant of the colonised site in accordance with the assumption of local ecological saturation.

The steady state OFD for this process is such that, in a large metacommunity with a total of *Z* slots, the mean number of species occupying *n* slots is *Zs*_*n*_, where for *m* = 0

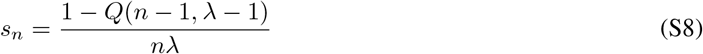

and for *m >* 0

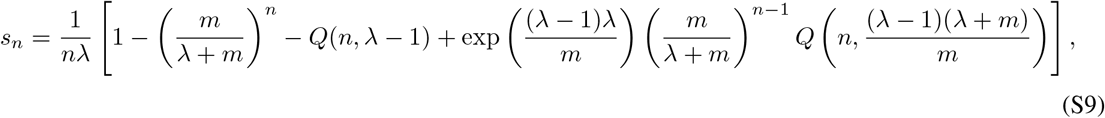

with *Q*(*a, x*) = Γ(*a, z*)*/*Γ(*z*) denoting the regularised incomplete gamma function. Noting that the total number of species in this metacommunity is *ZΣ*_*n*_*s*_*n*_, the definition of *λ* translates to

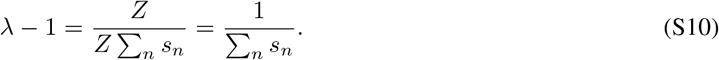

This is solved numerically by adjusting *λ*. In Fig. S3 we show the average occupancy frequency of the steady state of 1000 spatially implicit patch dynamics models assembled according to the basic principles described above. The dashed line represents the predicted OFD after numerically fitting the mixing rate.

To find the best fitting abiotic niche width *w* for an empirical dataset, we estimated the mixing rate *m* for the empirical OFD, then, for model metacommunities with abiotic variables extracted from observation data as described above, we scanned assembled metacommunities with various combinations of the focal parameters (Fig. S4). The best fitting *w* was selected as that with emergent OFD which closest matched the mixing rate *m* of the observation. Note that, for simplicity we define the site occupancy with reference to source populations only. These are the local populations which persist following a long relaxation (10^4^ unit times) after dispersal is switched off. In effect this assumes that only source populations are detected in the empirical data due to type II sampling errors. Alternative approaches to defining local site occupancy in simulations are available but typically require additional model assumptions or setting an arbitrary numerical threshold.

**Figure S3:**
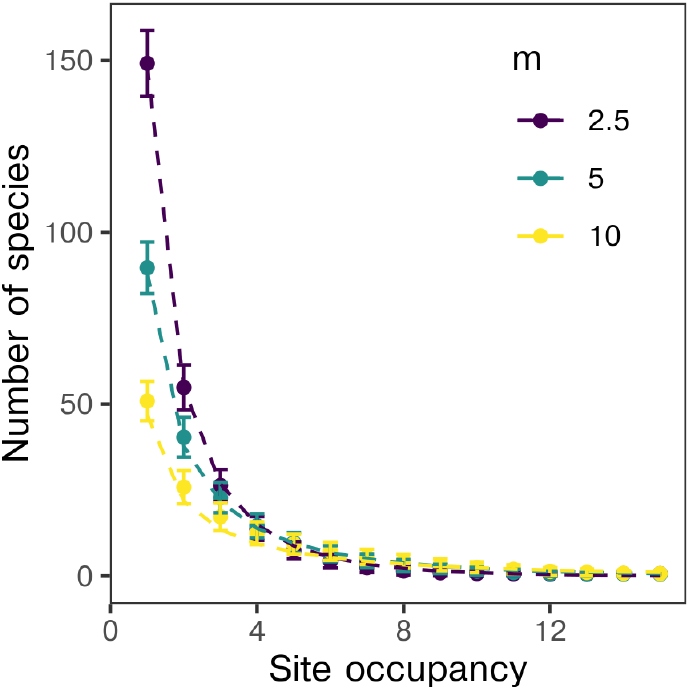
The steady state OFD of the locally saturated patch dynamics model. Points represent the average number of species in each site occupancy for 1000 realisations of the LSPOM described in the text for three different values of the mixing rate *m*. Dashed lines are the predicted OFDs computed by numerically solving Eq. (S10). Figure taken from O’Sullivan et al. (2021b).

**Figure S4:**
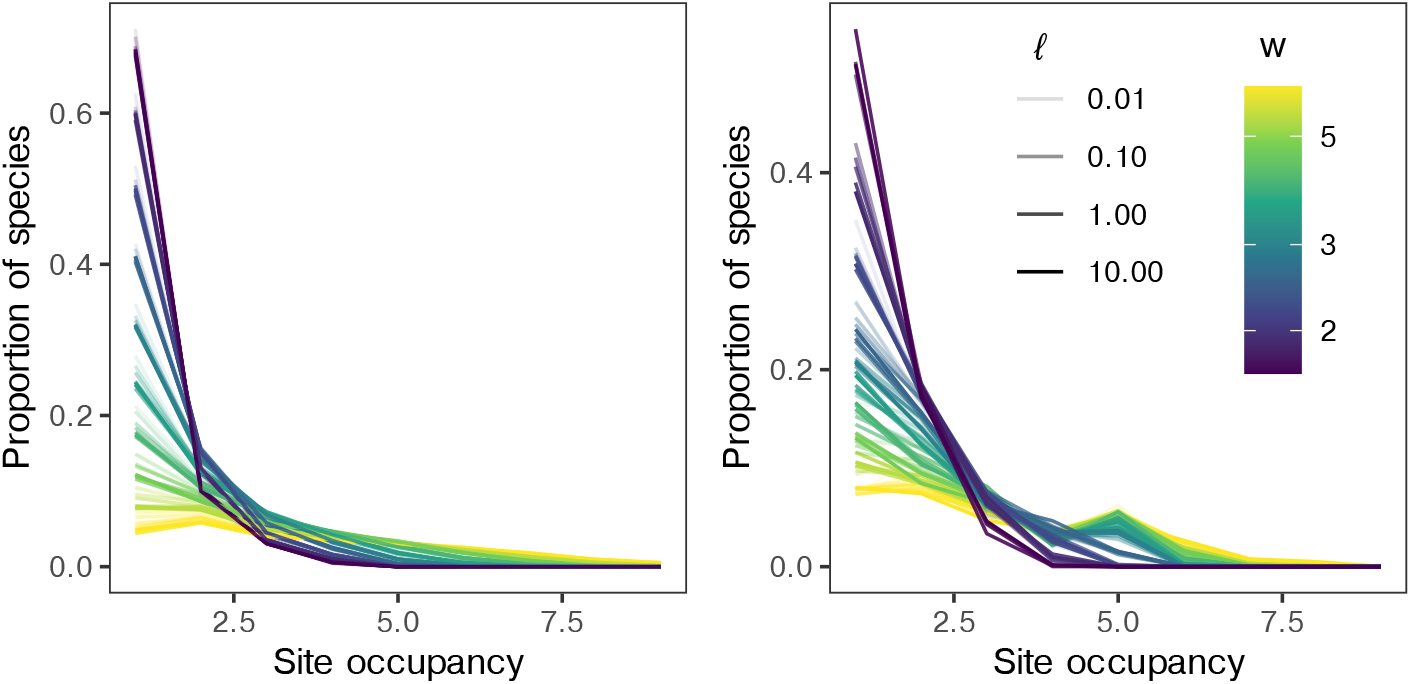
The impact of parameter variation on the OFD for fitted models. The average proportional occupancy frequency distributions for various values of *w* with landscape properties extracted from empirical datasets L and D. Colour represents abiotic niche width *w*.

### S4 Remote sensing data collection

We extracted temperature, precipitation and elevation data for the 23 sites of the Brazilian Lichen-Fungi survey (da Silva Cáceres et al., 2017). The first two abiotic variable were time averaged over the 20 years from January 2000 to January 2020. We pooled nearby sites in order to reduce the dimensionality and therefore the computational load of the simulations. For these sites we took abiotic values, including spatial coordinates, as the between site averages.

Temperature data were collected from the MODIS satellite database (Hulley and Hook, 2017). For each site, sensor products were filtered by land surface temperature and emissivity to retrieve data from the MODIS thermal infrared bands. Precipitation data were collected from the IMERG satellite database (Huffman et al., 2014). Elevation data were taken from the GTOPO30 digital elevation model database (Danielson and Gesch, 2011) which provides data from collective satellite inputs, compiled by the U. S. Geological Survey.

## Footnotes

1 CBD, https://www.cbd.int/doc/c/abb5/591f/2e46096d3f0330b08ce87a45/wg2020-03-03-en.pdf

1 https://www.cbd.int/sbstta/sbstta-24/post2020-indicators-en.pdf

